# An amino acid polymorphism in the *Drosophila* insulin receptor demonstrates pleiotropic and adaptive function in life history traits

**DOI:** 10.1101/008193

**Authors:** Annalise B. Paaby, Alan O. Bergland, Emily L. Behrman, Paul S. Schmidt

**Affiliations:** Department of Biology, University of Pennsylvania, Philadelphia, PA, 19104; Current address: Center for Genomics & Systems Biology, Department of Biology, New York University, New York, NY, 10003; Department of Biology, Stanford University, Stanford, CA, 94305

## Abstract

Finding the specific nucleotides that underlie adaptive variation is a major goal in evolutionary biology, but polygenic traits pose a challenge because the complex genotype-phenotype relationship can obscure the effects of individual alleles. However, natural selection working in large wild populations can shift allele frequencies and indicate functional regions of the genome. Previously, we showed that the two most common alleles of a complex amino acid insertion-deletion polymorphism in the *Drosophila* insulin receptor show independent, parallel clines in frequency across the North American and Australian continents. Here, we report that the cline is stable over at least a five-year period and that the polymorphism also demonstrates temporal shifts in allele frequency concurrent with seasonal change. We tested the alleles for effects on levels of insulin signaling, fecundity, development time, body size, stress tolerance, and lifespan. We find that the alleles are associated with predictable differences in these traits, consistent with patterns of *Drosophila* life history variation across geography that likely reflect adaptation to the heterogeneous climatic environment. These results implicate insulin signaling as a major mediator of life history adaptation in *Drosophila*, and suggest that life history tradeoffs can be explained by extensive pleiotropy at a single locus.

## INTRODUCTION

Many organisms display intraspecific variation in life history traits, including differences in reproductive timing and allocation, development, body size, stress tolerance, and lifespan. Alternative life history strategies have long been hypothesized to represent adaptive responses to variable selection (Stearns 1992), but examples with known causal nucleotides are rare (Flatt & Heyland 2011). However, sequence polymorphisms have been identified recently for flowering time in *Arabidopsis thaliana* (Méndez-Vigo et al. 2013) and adaptation to freshwater environments by marine stickleback fish, which includes changes in life history as well as in physiology and morphology (Jones et al. 2012). One reason that identification of molecular targets of selection is valuable is because it can illuminate mechanisms of multi-trait correlation. For example, the gene *Catsup* in *Drosophila melanogaster* affects longevity, locomotor behavior, and bristle number, but individual polymorphisms within the locus act on the traits independently and do not show pleiotropic effects (Carbone et al. 2006). Likewise, coat color in deer mice is a multi-phenotype trait associated with the *Agouti* locus, but mutations within the gene appear to have been targeted independently by selection with minimal pleiotropy (Linnen et al. 2013). These examples suggest that fitness-related traits, even those affected by the same gene, may be functionally independent such that natural populations might achieve any genetic and phenotypic combination. Alternatively, life history traits routinely exhibit strong genetic correlations, including negative associations between lifespan and reproduction and positive associations between lifespan and stress resistance (Reznick 1985; Stearns 1991; Partridge et al. 2005; Vermeulen & Loeschcke 2006; Harshman & Zera 2007; Toivonen & Partridge 2009). These associations mirror pleiotropic effects of lab-derived mutations and can be difficult to break in artificial selection experiments (Leroi et al. 2005; Patridge et al. 1999; Anderson et al. 2011; but see Khazaeli & Curtsinger 2010; Khazaeli & Curtsinger 2013). Consequently, pleiotropy, and specifically antagonistic pleiotropy, in which a genetic element encodes both positive- and negative-fitness phenotypes, remains an important component in the discussion of how life histories evolve (Williams 1957; Flatt & Heyland 2011).

*D. melanogaster* are distributed across environments that range from temperate to tropical, and exhibit variation in life history that appears adaptive. At high latitudes, populations exhibit higher incidence of reproductive diapause, larger body size, higher cold stress tolerance, and longer lifespan, but lower fecundity, relative to low latitude populations (Capy et al. 1993; Mitrovski & Hoffmann 2001; de Jong & Bochdanovits 2003; Schmidt et al. 2005a; Trotta et al. 2006). There is substantial genetic variance for these traits and pervasive genetic correlations among them, indicating that selection in the local environment may act on some phenotypes but drive expression of others through tradeoffs (David 1975; Anderson et al. 2003; de Jong & Bochdanovits 2003; Schmidt et al. 2005b; Rako et al. 2007; Schmidt & Paaby 2008). Tolerance to environmental stress may be especially important, and the ability to resist desiccation, starvation and temperature stress correlates with climatic environment (Hoffmann & Harshman 1999; Hoffmann et al. 2001; Hoffmann et al. 2005; Hoffmann et al. 2007). This framework suggests a hypothetical selection regime: at high latitude, cold winters impose seasonal stress and favor genotypes that confer stress tolerance and overwintering ability; correlated traits such as larger body size, longer lifespan, slower development and lower fecundity may evolve as co-adapted responses to the same selection regime or by indirect selection via pleiotropy (Paaby & Schmidt 2009).

Few loci have been found to explain the observed genetic variance for *D. melanogaster* life history (De Luca et al. 2003; Carbone et al. 2006; Paaby & Schmidt 2008; Schmidt et al. 2008; Bergland et al. 2012; Remolina et al. 2012, Sgrò et al. 2013). Genes in the Insulin/Insulin-like Signaling (IIS) pathway are good candidates because experimental manipulation of IIS mirrors the life history tradeoffs observed in natural populations (Partridge & Gems 2002; Tatar et al. 2003; Giannakou & Partridge 2007; Grönke 2010). We previously evaluated sequence variation at the *Insulin-like Receptor* (*InR*) in wild populations of *D. melanogaster* and observed striking reciprocal clines in the northern and southern hemispheres for the common alleles of a complex amino acid indel polymorphism (Paaby et al. 2010). The polymorphism disrupts a region of glutamine-histidine repeats in the first exon (Guirao-Rico & Aguadé 2009). In both North America and Australia, the most common allele is at low frequency in tropical and subtropical populations and increases in frequency with latitude, showing highest frequency in temperate populations; the second most common allele exhibits the inverse cline. These patterns appear to be responses to similar but independent selection pressures between the continents: fly populations in North America and Australia were founded at different times and from different source populations (Bock & Parsons 1981; David & Capy 1988), and nucleotide variants on either side of the polymorphism show neutral patterns across geography.

Initial tests showed that the two common alleles, which differ in length by two amino acids and are herein designated *InR^short^* and *InR^long^*, demonstrated significant effects on fitness traits. *InR^short^*, the allele common at high latitudes, was associated with greater stress tolerance; *InR^long^*, which is common at low latitudes, was associated with greater fecundity (Paaby et al. 2010). Here we show that the allele frequency cline in North America persists five years later and that temporal changes in allele frequency within a single population correlate with fluctuations in the seasonal environment, which mimic the temperature and resource availability differences associated with latitudinal climate. We also find that the *InR* locus in general exhibits elevated patterns of clinality and seasonality relative to the rest of the genome. We more thoroughly tested the functional effects of the complex polymorphism by measuring levels of IIS and a spectrum of life history phenotypes in *InR^short^* and *InR^long^* lines with randomized genetic backgrounds. We find that the alleles associate predictably with the tested traits, suggesting that the complex amino acid indel polymorphism at *InR* is an important target of contemporaneous selection in wild *D. melanogaster* populations, is highly pleiotropic, and contributes to observed tradeoffs in *D. melanogaster* life history.

## METHODS

***Allele frequency estimates***. As described in Bergland et al. (In Review), between 50-200 isofemale lines were established from six populations spanning Florida to Maine along the North American east coast (latitudes 25.5°N, 30.9°N, 33.0°N, 35.5°N, 39.9°N, 44.1°N). The population from Linvilla Orchard in Media, PA (39.9°N, 74.4°W) was additionally sampled in the spring and fall in 2009, 2010, and 2011. Within five generations a single male from each line was pooled into population samples, which were sequenced on an Illumina HiSeq2000 using 100-bp paired-end reads. Reads were aligned to the *D. melanogaster* reference genome v5.39 using *bwa* v0.5.9-r16 (Li & Durbin 2009). The indel polymorphism at *InR* was called using *UnifiedGenotyper* from GATK v3.1-1 (McKenna et al. 2010), which identified multiple discrete polymorphisms within the complex indel haplotypes that we previously characterized (Paaby et al. 2010). That characterization identified many haplotypes of different lengths, two of which were common (herein *InR^short^* and *InR^long^*); the reference genome to which the new pooled samples were mapped has a haplotype of intermediate length (previously referred to as *InR^251^*). Thus, the complex haplotypes, previously identified by their lengths, were decomposed into sets of shared or distinct alleles at discrete polymorphic sites.

We used our prior Sanger sequence data (see GenBank accessions GQ927177–GQ927244) to assign alleles of the discrete polymorphisms to the *InR^short^* and *InR^long^* allele classes. Here we report those classifications (polymorphisms are denoted by their position in the *D. melanogaster* reference genome v5.48). Three discrete polymorphisms in the pooled sequencing data discriminate unambiguously between *InR^short^* and *InR^long^*: 17405631-7, 17405651-4 and 17405784. In each case, the reference genome allele matches *InR^short^*. Only one of the discrete polymorphisms, SNP 17405614, was variable within an allele class; the alternate and reference alleles are both associated with *InR^short^* such that the association between the discrete allele identities and the larger haplotypes in which they are embedded (*InR^short^* and *InR^long^*) are approaching equilibrium (R^2^=0.45). SNP 17405619 segregates at low frequency and was invariant in our Sanger sequencing data such that the *InR^short^* and *InR^long^* haplotypes we observed shared the same discrete allele. Both the alternate allele of SNP 17405631 and the alternate allele of 17405634-7 match both *InR^short^* and *InR^long^*. For polymorphism 17405637, two alternate alleles were called; the 6-nt insertion matches *InR^long^*, but the 3-nt insertion is confounded with the alternate allele of 17405634-7 (which is called consistently). Thus, we ignored the 3-nt insertion allele at 17405637.

Allele frequencies of SNPs across the entire *InR* locus and across other sites on chromosome 3R were called with CRISP (Bansal 2010) as described in Bergland et al. (In Review). To test whether the *lnR* locus (3R:17385970-17455043) demonstrates elevated clinality or seasonality relative to the rest of the genome, we calculated the proportion of SNPs within *InR* that were identified to be clinal (with associated clinal FDR < 0.01) or to vary repeatedly among seasons over the course of three years (with associated seasonal FDR < 0.5) using estimates of clinality and seasonality reported in Bergland *et al* (In Review). We note that the high FDR for seasonal SNPs was chosen to increase power for detecting fluctuations at *InR* outside of the *InR^long/short^* polymorphism, which is not included in this scan or in the dataset evaluated by Bergland et al. (In Review). We contrasted the proportion of clinal and seasonal SNPs within the *InR* locus to 1000 random chromosomal regions matched for length, chromosome, and inversion status for *In(3R)Payne*.

***Genotyping***. To genotype flies for the experimental assays, the identity of the complex *InR* indel was determined by measuring its amplified fragment length following polymerase chain reaction (PCR) with a fluorescent-tagged primer on an Applied Biosystems 3100 capillary sequencer. PCR conditions, including primer sequences, are described in Paaby et al. (2010). That paper reported the distribution of alleles across geography and referred to the alleles by their PCR fragment lengths, which we erroneously stated as 248 bases for the high-latitude allele and 254 bases for the low-latitude allele. The actual fragment lengths using the cited primers are 5 bases longer. To avoid confusion, in this paper we refer to these two alleles as *InR^short^* (previously *InR^248^*) and *InR^long^* (previously *InR^254^*). It should be noted that although fragment-length genotyping does not discriminate between the SNP alleles we characterized in or near the complex indel polymorphism (described above), only one of those SNPs (17405614) appears to be independent of the *InR^short^* and *InR^long^* allele classes. All evidence to date indicates that fragment-length genotyping unambiguously discriminates between consistent, but complex, haplotype classes (see Figure 1A-B).

**FIGURE 1.**
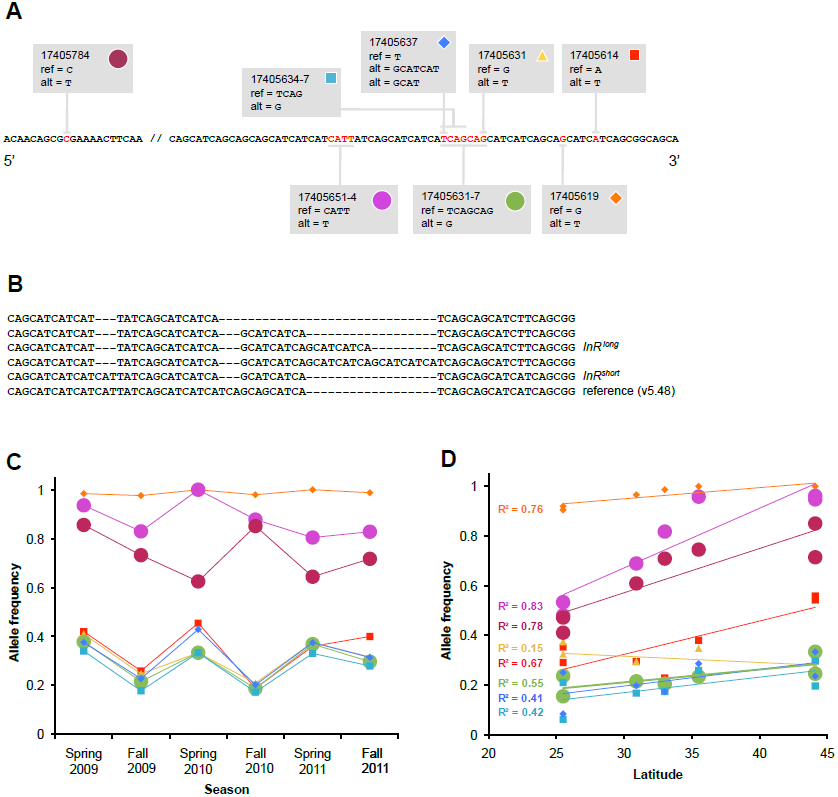
From pooled sequencing data, we identified eight discrete polymorphisms associated with the complex indel in the first exon of *InR* (A). Each polymorphism is labeled according to position on chromosome 3R relative to the published *D. melanogaster* reference genome (v5.48). The discrete polymorphisms describe the complex indel haplotypes we previously characterized, which include two alleles, *InR^short^* and *InR*long**, that are common at high and low latitudes respectively (B). Polymorphisms that unambiguously discriminate between the *InR^short^* and *InR^long^* alleles are denoted with a large dot; in each case, the allele present in the reference genome identifies *InR^short^*. (See Materials & Methods for relationships between the other discrete polymorphisms and *InR^short^* and *InR^long^*.) All eight of the discrete polymorphisms demonstrate some degree of seasonality, but four exhibit especially strong patterns in which the plotted allele increases in frequency by approximately 20% over the winter for three consecutive years (C). Alleles assigned to the *InR^short^* allele class (large dots) generally show higher frequency in the spring and lower frequency in the fall, suggesting that overwintering may impose selection pressures consistent with high latitudes. As in our earlier report (Paaby et al. 2010), alleles assigned to the *InR^short^* allele class also increase in frequency with latitude (D). The plotted alleles are all reference genome alleles.

***Fly stocks***. To determine whether *InR^short^* and *InR^long^* alleles have different effects on phenotype, we generated stocks with *InR^short^* or *InR^long^* on the third chromosome, and for which the X and second chromosomes were replaced so that they were isogenic within and across lines. The genome regions surrounding *InR* on the third chromosome were randomized, from lines originally genotyped as *InR^short^* and *InR^long^*, across lines. To generate the stocks, we selected *InR^short^* and *InR^long^* parental strains, neither of which carried the *In(3R)Payne* inversion, and for which the X and second chromosome were already replaced (for the X chromosome, stock 2475; for the second chromosome, stock 6326 from the Bloomington Stock Center; originally described in Paaby et al. 2010). The parental strains were crossed and the offspring permitted to recombine for four generations, so that genetically variable regions from the parental third chromosomes were distributed as randomized blocks across lines. Fourteen isogenic stocks were subsequently established per allele by homozygosing the third chromosome via balancer extraction and genotyping the *InR* indel polymorphism. This entire scheme was performed twice, using lines derived from two independent populations: Mount Sinai, NY (40.95°N, 72.84°W) and Bowdoinham, ME (44.01°N latitude, 69.90°W longitude).

*InR^short^* and *InR^long^* alleles were evaluated as homozygotes and heterozygotes, as well as with *InR* mutant and wild-type alleles derived from a laboratory strain. The *InR* mutant was the hypomorphic *InR^p5545^* allele generated by P-element insertion in exon 1 (Tatar et al. 2001) balanced over the TM3 chromosome (stock 11661 from the Bloomington Stock Center, ry^506^ P{PZ}InR^p5545^/TM3, ry^RK^ Sb^1^ Ser^1^). The TM3 balancer contains a wild-type *InR* allele (herein *InR^TM3^*) that is intermediate in length between *InR^short^* and *InR^long^*: it is three nucleotides longer than *InR^short^*, corresponding to an additional histidine, and three nucleotides shorter than *InR^long^*, which contains an additional glutamine.

***Flies for assays***. Once the stocks were established, we used three methods to generate flies for the phenotype assays (derived from one or both of the independent populations). The first two tested the effects of *InR^short^* and *InR^long^* in a wild-type background; the third tested *InR^short^* and *InR^long^* effects while paired with either the *InR* hypomorphic allele or a wild-type allele in the TM3 balancer chromosome. In the first method, we exerted nominal control over the background variation in the stocks by crossing 14 *InR^short^* and 14 *InR^long^* stocks in a round-robin design to generate a total of 42 lines from which we collected flies for phenotyping (14 each of *InR^short^/InR^short^*, *InR^long^/InR^long^*and *InR^short^/InR^long^*). The crosses were conducted in standard media vials with 7 virgin females (2-5 days old) and 4 young males (3-8 days old). This design produced non-independent replication within each genotypic class, and because test stocks were never mated to themselves, also ensured that no individuals were strictly isogenic at the third chromosome. In the second method, we combined stocks (minimum of five) carrying the same allele (*InR^short^* or *InR^long^*) but otherwise randomized variation on the third chromosome in bottles, permitting the stocks to continue recombining freely. To rear flies for the assays, 40 virgin females (2-5 days old) and 20 young males (3-8 days old) were collected and mated in fresh bottles, either within or across genotype class, to generate the two homozygous and the heterozygous genotypes. Unless otherwise specified, the phenotype assays under this method were replicated via 10 mating bottles for each genotypic class. In the third method, we crossed 12 *InR^short^* stocks and 12 *InR^long^* stocks to the hypomorphic *InR^p5545^/TM3* stock (in vials as in method one). From these 24 lines, we collected progeny carrying both the *InR* hypomorphic allele and the wild-type allele in the balancer chromosome for phenotyping.

***Phenotype assays***. For all assays, flies were reared and assays were performed on standard cornmeal-molasses media at room temperature and subject to ambient light cycles. Larval density was kept low to limit overcrowding. We tested *InR^short^* and *InR^long^* alleles derived from Mount Sinai in all assays; we tested *InR^short^* and *InR^long^* alleles from Bowdoinham in a subset of the assays. All *InR^short^* and *InR^long^* alleles tested in the presence of the *InR* hypomorph (*InR^p5545^*) or wild-type balancer (*InR^TM3^*) were from Mount Sinai. All statistics were performed using JMPv7 (SAS Institute, Cary, NC). Within each analysis, we used planned comparisons to test for significant differences between *InR^short^* and *InR^long^* genotypes.

*<u>Lifespan and lifetime fecundity</u>*. Bottle cages were populated with 30 virgin females and 30 males, collected over 24 hours, from lines generated by method one from the Mount Sinai population. A total of 42 bottles, comprising 14 of each genotypic class, were maintained by changing standard media egg-laying plates every day (days 1-26) or every other day (days 28 onward) until all flies were dead. Bottles were inverted, so that dead flies collected on the media plates. Dead flies and eggs laid were scored at every plate change. Lifespan was analyzed by a proportional hazards model, and cumulative fecundity was analyzed by ANOVA.

*<u>Early fecundity</u>*. Virgin females and males were collected over 6 hours, separated into single-sex vials of 5 females or 3 males each, and aged for 3 days on standard media with topical yeast. Flies were transferred in mixed-sex batches (five females and three males) to media-free vials and starved for 4 hours, then transferred again to fresh media vials and permitted to lay eggs. The number of eggs laid was recorded after 12 hours and analyzed by ANOVA. Flies were generated by method two from both populations, and also with the *InR* hypomorph via method three.

*<u>Development time</u>*. Young (3-6 days old) males and females (5 flies of each sex) were transferred from standard media vials to media-free vials and starved for four hours, then transferred again to fresh media vials and permitted to lay eggs, at densities of ≤30 eggs per vial, for up to four hours before being removed. Flies derived from Mount Sinai were generated via method one; flies from Bowdoinham via method two, and with the hypomorph via method three. The number of adults per vial was recorded daily at 8 AM, 12 PM and 8 PM for all emerging adults, and eclosion time was analyzed by ANOVA.

*<u>Body weight, lipid content and body size</u>*. Using methods and replication schemes identical to those described for development time, eggs were reared and permitted to eclose as adults. After eclosion, adults were transferred to fresh media vials containing topical yeast and aged for 24 hours, then frozen at −80°C. To measure dry weight, frozen flies were dried for 24 hours at 55°C and weighed to 0.0001 g in single-sex batches of 7 individuals on a Sartorius microbalance (1-2 batches per original vial cross). The batches were transferred to 1.5 mL tubes and lipids were extracted by adding 400 uL diethyl ether and gently rocking the tubes for 24 hours. Flies were dried overnight at 55°C and batches re-weighed; lipid content was computed by subtraction. To estimate body size, left wings were removed from frozen flies and affixed to an index card using clear tape. Pictures of wings from a DC 300 camera (Leica) on a dissecting microscope were measured for L2 and L3 vein length using landmarks 6, 12 and 13 (Gidaszewski et al. 2009) on 3-8 males and 3-8 females from each vial cross, using tpsDigv2.12 (Rohlf 2008). By the same method thorax width was measured for the Bowdoinham and hypomorph crosses. Measurements were analyzed by ANOVA.

*<u>Chill coma recovery</u>*. Freshly eclosed flies were collected over 24 hours from flies generated by method one from the Mount Sinai population, and sorted into standard media vials in batches of 5 males and 5 females each. After 3 days, males were sorted into fresh vials and aged another day. To induce chill coma, vials were covered in ice and placed at 4°C for three hours, then restored to room temperature. Time to recovery (transition to the upright position) was recorded using a video camera and analyzed by ANOVA.

*<u>Cold shock and starvation</u>*. Freshly eclosed flies were collected over 24 hours from flies generated by method one from the Mount Sinai population and method three with the hypomorph, sorted into standard media vials in batches of 5 males and 5 females, and aged for 3 days. The 42 vials generated by method one were replicated twice. To induce cold shock, vials were inverted and exposed to −20°C (air temperature) for 25 minutes. After 24 hours, individuals were scored as dead or alive. To measure starvation resistance, flies were transferred into media-free vials containing a cotton ball saturated with 2 mL of water. At 51 hours, males were scored as dead or alive; at 64 hours, females were scored as dead or alive. Both assays were analyzed by nominal logistic regression modeling the log odds of mortality/survivorship.

*<u>Heat shock</u>*. Freshly eclosed flies were collected over 24 hours and sorted into mixed-sex standard media vials of 5 males and 5 females each. Flies were generated by method two from the Mount Sinai population (15-20 replicate vials per genotypic class) and by method three. Flies were aged for 4 days and then transferred to fresh media vials. Vials were inverted, placed at 25°C, and warmed to 37°C; after reaching 37°C (approximately 20 minutes), flies were kept at 37°C for 25 minutes and then removed to room temperature to recover. After 24 hours, flies were scored as dead or alive and analyzed by nominal logistic regression modeling the log odds of mortality/survivorship.

***Quantitative PCR***. To test whether *InR* alleles affect IIS, we used quantitative polymerase chain reaction (qPCR) to determine the relative abundance of 7 transcriptional targets of dFOXO, a central transcription factor within the IIS pathway (Jünger et al. 2003; Puig et al. 2003; Wang et al. 2005; Casas-Tinto et al. 2007; Flatt et al. 2008; Vihervaara & Puig 2008; Matilla et al. 2009). Total RNA was prepared from flies generated by method two from flies derived from Mount Sinai. Thirty virgin females from each of the three genotypic classes were collected over 24 hours, aged on standard media, and snap frozen. RNA was extracted using RNeasy (Qiagen) and reverse transcribed using iScript cDNA synthesis kit (Bio-Rad), following the manufacturer's protocol. Relative abundance of transcript levels was determined using an ABI 7500 Fast Real-Time PCR machine and SYBR Green PCR Master Mix (Applied Biosystems) by the DDC_T_ relative quantitation method. Four technical replicates were used for each sample and relative abundance was normalized by using *GAPDH2* as an endogenous control. Other work has demonstrated that *GAPDH2* is an appropriate control for measuring relative levels of IIS (Hwangbo et al. 2004, Flatt et al. 2008). Primer sequences for the qPCR reaction are: *GAPDH2* forward, GCGGTAGAATGGGGTGAGAC; *GAPDH2* reverse, TGAAGAGCGAAAACAGTAGC; *4E-BP* forward, GAAGGTTGTCATCTCGGATC; *4E-BP* reverse, ATGAAAGCCCGCTCGTAGA; *l(2)efl* forward, AGGGACGATGTGACCGTGTC; *l(2)efl* reverse, CGAAGCAGACGCGTTTATCC; *ac76e* forward, CAGGATGAATGACGCCCTTTCGG; *ac76e* reverse, ATGGACACAACACATGCCAGCAGC; *dLip4* forward, GATAGCAATGTGCGGTTGGA; *dLip4* reverse, TCATCCGTCTCCAAGGTGTG; *InR T1* forward, CACAAGCTGGAAAGA AAGTGC; *InR T1* reverse, CAAACACGTTTCGATAATATTTTTCT; *InR T2* forward, GCCTCGCACTTTGCTTATGT; *InR T2* reverse, AAAAACAACGACAGCGACAA; *InR T3* forward, TTACGCCACTGCATTCGTTC; *InR T3* reverse, ATGGCCTCTCTCTCCGTCTC.

## RESULTS

***Clinal and seasonal estimates of allele frequency*.** To test whether *InR* allele frequencies change in association with the climatic environment, we identified multiple discrete polymorphisms within and near the complex indel polymorphism from pooled population samples. We identified all variants (SNPs or small indels) that comprise the complex indel itself and three SNPs −11, −16 and +130 nucleotides from the indel that segregate at frequencies above 5% (Figure 1A-B). Four of these discrete polymorphisms show clear seasonality, with repeating fluctuations in allele frequency of approximately 20% between spring and fall time-points over three consecutive years (Figure 1C). The other four discrete polymorphisms are suggestive of seasonality: they demonstrate either comparable changes in amplitude but a deviation in the frequency pattern at a single time-point, or a repetitive pattern at low amplitude. The discrete polymorphisms also show clines in allele frequency across latitude (Figure 1D). We were able to unambiguously assign allele identities for three of the discrete polymorphisms to the *InR^short^* or *InR^long^* allele classes (Table S1), which exhibited strong allele frequency clines five years previously (Paaby et al. 2010). In each of these cases, changes in allele frequency across latitude replicated our earlier observations, and seasonal fluctuations in frequency for these discrete polymorphisms support the hypothesis that alleles favored at high latitude are favored during the winter.

**TABLE 1.** Statistical results for tests of fecundity, development, body size and chill coma recovery for *InR* genotypes in a wild-type background.

|  | *Mount Sinai population* |  |  |  |  |  | *Bowdoinham population* |  |  |  |  |
| --- | --- | --- | --- | --- | --- | --- | --- | --- | --- | --- | --- |
|  | Source | DF | SS | F | p |  | Source | DF | SS | F | p |
| Lifetime fecundity | genotype | 2 | 6160.87 | 2.4026 | 0.1038 |  |  |  |  |  |  |
| Early fecundity | genotype | 2 | 1469.05 | 55.85 | <0.0001 |  | genotype | 2 | 335.23 | 8.02 | 0.0018 |
| Development time | genotype | 2 | 81440.13 | 270.48 | <0.0001 |  | genotype | 2 | 2742.59 | 1.24 | 0.2891 |
|  | sex | 1 | 271.65 | 1.81 | 0.1793 |  | sex | 1 | 2449.16 | 2.22 | 0.1367 |
|  | genotype x sex | 2 | 245.64 | 0.82 | 0.4424 |  | genotype x sex | 2 | 9928.1 | 4.5 | 0.0115 |
| Dry body weight | genotype (f) | 2 | 2.57x10<sup>-6</sup> | 19.51 | <0.0001 |  | genotype (f) | 2 | 2.21x10<sup>-6</sup> | 80.95 | <0.0001 |
|  | genotype (m) | 2 | 4.14x10<sup>-7</sup> | 14 | <0.0001 |  | genotype (m) | 2 | 4.28x10<sup>-7</sup> | 33.52 | <0.0001 |
| Lipid weight | genotype | 2 | 0.1242 | 3.7185 | 0.0282 |  | genotype | 2 | 0.4206 | 19.1552 | <0.0001 |
|  | sex | 1 | 0.1532 | 9.1744 | 0.0032 |  | sex | 1 | 0.3181 | 28.9744 | <0.0001 |
|  | genotype x sex | 2 | 0.0161 | 0.4833 | 0.6184 |  | genotype x sex | 2 | 0.077 | 3.5062 | 0.035 |
|  | mass | 1 | 0.022 | 1.3187 | 0.2539 |  | mass | 1 | 0.1656 | 15.083 | 0.0002 |
| L2 wing vein | genotype (f) | 2 | 0.0253 | 7.3131 | 0.0008 |  | genotype (f) | 2 | 0.5059 | 189.483 | <0.0001 |
|  | genotype (m) | 2 | 0.0153 | 4.1532 | 0.017 |  | genotype (m) | 2 | 0.3939 | 21.5208 | <0.0001 |
| Wing:thorax ratio |  |  |  |  |  |  | genotype (f) | 2 | 0.0627 | 18.7883 | <0.0001 |
|  |  |  |  |  |  |  | genotype (m) | 2 | 0.0628 | 2.7051 | 0.0708 |
| Chill coma assay | genotype (f) | 2 | 1262285.3 | 5.49 | 0.0048 |  |  |  |  |  |  |
|  | genotype (m) | 2 | 829073.78 | 12.66 | <0.0001 |  |  |  |  |  |  |

Compared to the rest of the genome, the *InR* locus exhibits elevated patterns of allele frequency change across latitude and seasonal time. The region surrounding *InR* contains significantly more clinal polymorphisms relative to other regions of chromosome 3R matched for size and inversion status (42% vs. 20%, respectively; Pr(%InR > %control) = 0.986). Relative to matched control regions, the *InR* locus also contains slightly more seasonal polymorphisms (6% vs. 4.7%; Pr(%InR > %control) = 0.8). Previous examination of linkage disequilibrium across *InR* (Paaby et al. 2010) indicates that the complex indel polymorphism is largely independent of variation elsewhere in the gene, which implies that many sites within *InR* may be targets of spatially and temporally varying selection.

***Allele effects*.** We evaluated the effects of *InR* alleles on levels of IIS and multiple phenotypes and found that individuals carrying the *InR^short^* allele, which is which is prevalent in high-latitude, temperate environments, exhibited lower levels of IIS, better survived cold temperature stresses and starvation, and showed sex-specific evidence for longer lifespan. Flies carrying the *InR^long^* allele, which is common in low latitude, warm environments, were associated with increased signaling and exhibited higher rates of fecundity, larger body mass, faster development time and better survival of heat shock. These phenotypic differences are consistent with experimental reduction of IIS as well as life-history tradeoffs observed in natural populations, and suggest that a single locus can act pleiotropically on a suite of fitness-related traits. We replicated some of the assays by using *InR^short^* and *InR^long^* alleles derived from two independent populations (Bowdoinham and Mount Sinai); the results are reported separately.

***Levels of insulin signaling.*** To test whether the observed phenotypic differences between *InR^short^* and *InR^long^* are mediated by different levels of IIS, we used qPCR to measure relative abundance of seven transcriptional targets of dFOXO, a central transcription factor in the IIS pathway that is repressed by InR activity. Decreased InR activity increases target abundance, and transcripts for five of these (*4E-BP*, *l(2)efl*, *ac76e*, *dLip4* and *InR T1*) were highest in *InR^short^*, intermediate in the heterozygote, and lowest in *InR^long^*; all but *4E-BP* showed statistical significance (p<0.05) between *InR^short^* and *InR^long^* (Figure 2). This pattern is robust considering that the *InR^short^* genotype exhibited at most twofold higher levels relative to *InR^long^*: a difference that may be biologically meaningful, and realistic for wild-type genotypes, but close to the limit of detection by qPCR methods. Two other targets, transcripts of *InR* itself, show no difference in abundance level between the homozygote genotypes, but significantly higher abundance in the heterozygote.

**FIGURE 2.**
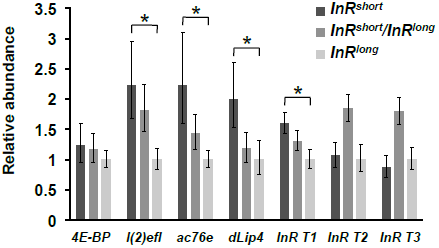
Relative abundance of seven transcriptional targets of dFOXO, a central transcription factor in the IIS pathway that is repressed by InR activity. Increased abundance of these targets indicates reduced insulin signaling; transcript abundance for *l(2)eft*, *ac76e*, *dLip4* and *InR T1* were significantly greater (p<0.05) in *InR^short^* samples compared to *InR^long^*. These results are consistent with the expectation that *InR^short^* is associated lower IIS than *InR^long^*. Transcripts were measured from flies derived from the Mount Sinai population. Error bars show 95% confidence intervals.

***Fecundity.*** Surprisingly, the homozygous *InR* genotypes showed no significant differences in total fecundity: a fly carrying the *InR^long^* allele laid on average only 5.3 more eggs over her lifetime than a fly carrying *InR^short^* (p=0.6949, Table 1). This was unexpected because we routinely observed substantially greater population numbers in our *InR^long^* bottle cultures, and previous results showed that the *InR^long^* genotype is more fecund (Paaby et al. 2010). Examination of the eggs laid per day, however, revealed that the *InR^long^* genotype lays more eggs early in life (Figure 3A). We performed another assay to explicitly measure early fecundity, or how many eggs young mated females laid upon first access to fresh food. Here, we found that the *InR^long^* genotype is nearly six times more fecund than *InR^short^* in the first 12 hours of egg-laying (p<0.0001, Table 1, Figure 3B). This result was again observed in a replicate assay using flies derived from the second population (p=0.0005) (Table 1, Figure 3C).

**FIGURE 3.**
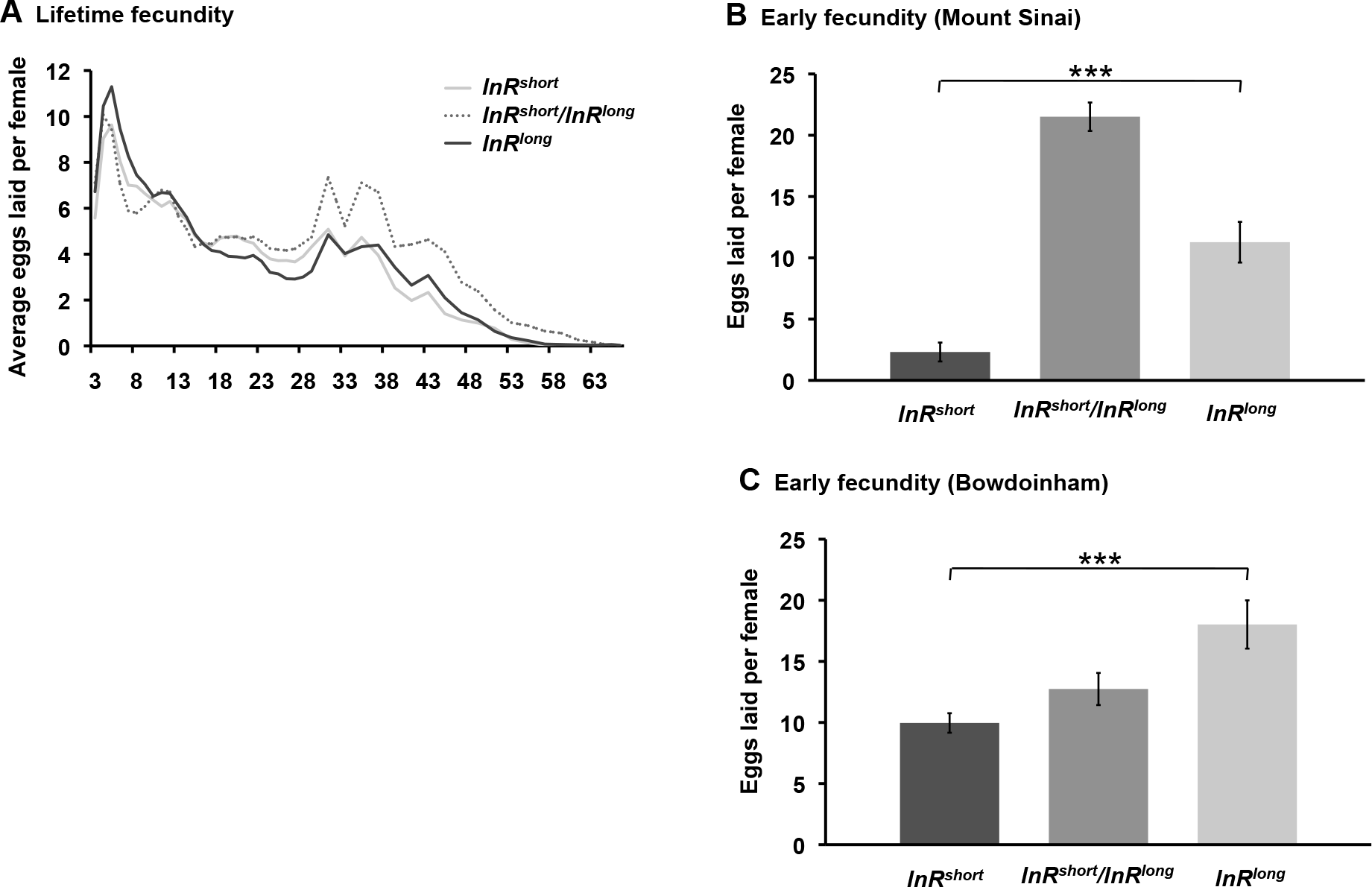
Average eggs laid per female over lifetime and in the first 12 hours after mating. Lifetime fecundity was measured in flies derived from the Mount Sinai population and was not significantly different between any genotypes except between *InR^short^* and the heterozygote (A). However, *InR^long^* females laid many more eggs in the first 12 hours after mating than *InR^short^* females, in comparisons from both Mount Sinai (DF=1, 21; F=22.79; p=0.0001) and Bowdoinham (DF=1, 27; F=15.55; p=0.0005) (C). Error bars show 95% confidence intervals.

***Development time.*** From egg to emerging adult, flies carrying the *InR^long^* allele developed faster than flies carrying *InR^short^*. This result was replicated in flies derived from two populations: in the first, males and females carrying *InR^long^* developed an average of 6.0 hours faster than those carrying *InR^short^* (p<0.0001); in the second population, development time was not significantly different among male genotypes, but *InR^long^* females emerged an average of 18.2 hours ahead of *InR^short^* females (p=0.0029) (Table 1, Figure 4).

**FIGURE 4.**
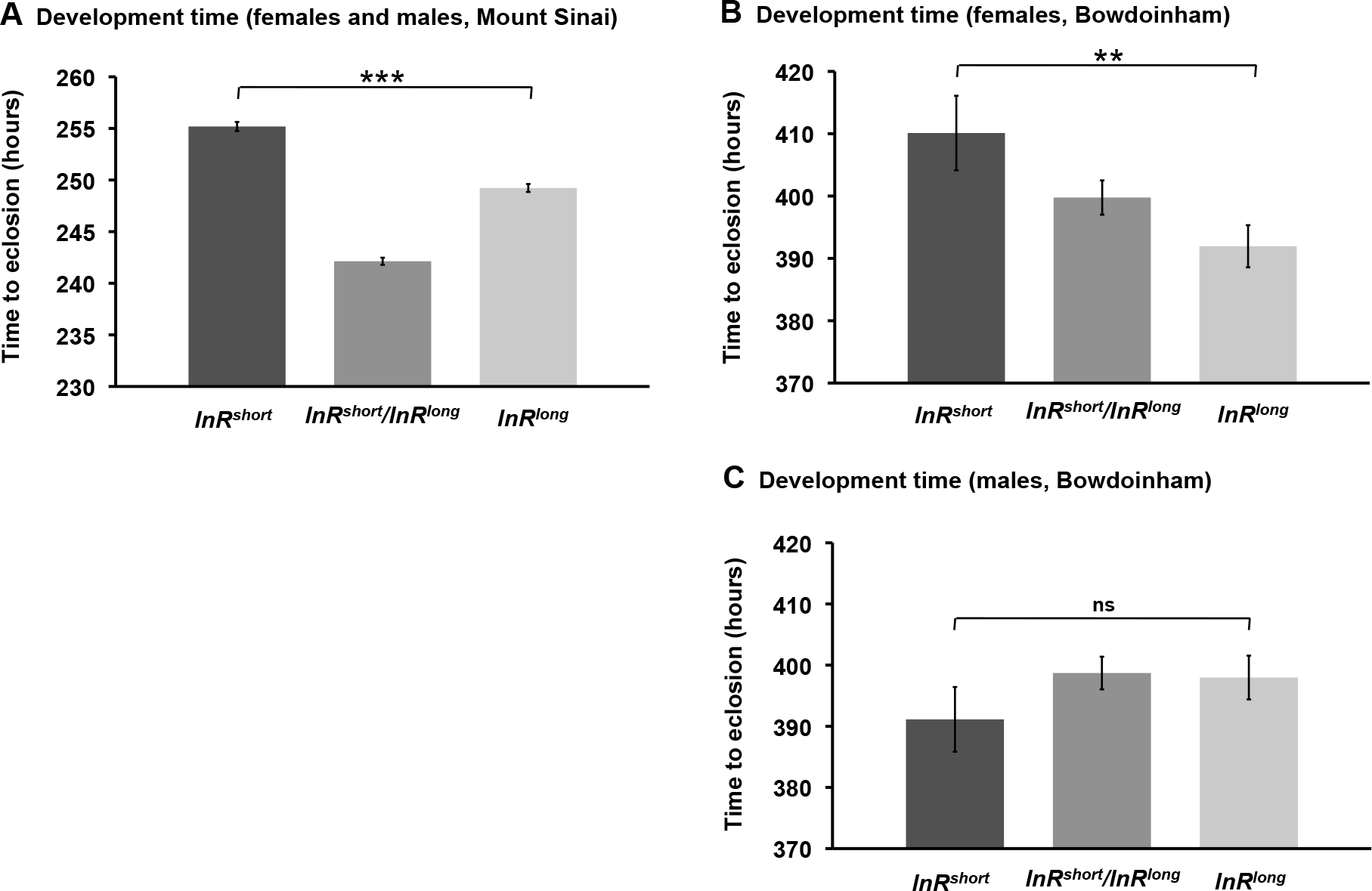
Average development time, from the time the egg was laid to eclosion of the adult. Flies derived from the Mount Sinai population showed no significant differences in development time by sex, and *InR^long^* flies developed faster than *InR^short^* flies (DF=1, 3006; F=113.23; p<0.0001) (A). Flies derived from Bowdoinham showed a significant sex effect; *InR^long^* females developed faster than *InR^short^* females (DF=1, 244; F=9.07; p=0.0029) (B), but male genotypes were not significantly different (DF=1, 278; F=1.42; p=0.2341) (C). Error bars show 95% confidence intervals.

***Body weight and size.*** Because reduction in IIS reduces body size (Clancy et al. 2001; Tatar et al. 2001), we predicted that individuals carrying the *InR^long^* allele would be larger than individuals carrying *InR^short^*. Indeed, the average dry body weights of *InR^long^* flies were heavier than *InR^short^* flies. This was true for alleles tested from both populations: *InR^long^* females and males derived from Bowdoinham were 26.1% and 18.6% heavier, respectively (p<0.0001 for both), while *InR^long^* females and males derived from Mount Sinai were 15.5% and 10.9% heavier (p=0.0002 and p=0.0004) (Table 1, Figures 5, S1). The higher mass associated with the *InR^long^* genotype is partially explained by lipid content. For both populations, the

**FIGURE 5.**
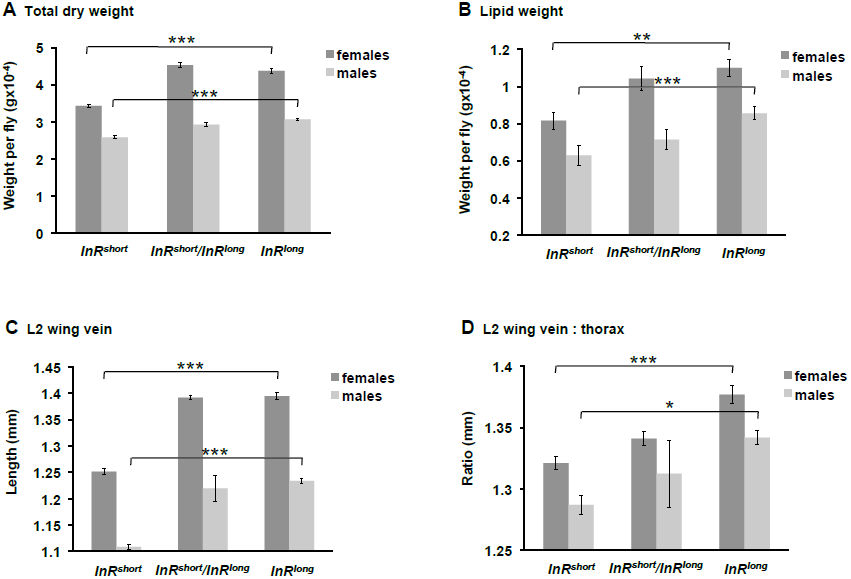
Average dry body weight, lipid weight, length of the L2 wing vein, and wing:thorax ratio for flies derived from the Bowdoinham population. *InR^short^* flies had lower dry weight (A) and lower lipid content (B) than *InR^long^* flies for both females (DF=1, 37; F=103.96; p<0.0001 for dry weight; DF=1, 36; F=11.09; p=0.002 for lipid weight) and males (DF=1, 39; F=63.17; p<0.0001 for dry weight; DF=1, 38; F=27.36; p<0.0001 for lipid weight). *InR^short^* flies were also smaller, as measured by the L2 wing vein (DF=1, 117; F=297.32; p<0.0001 for females; DF=1, 123; F=35.86; p<0.0001 for males) (C), but had a smaller wing:thorax ratio than *InR^long^* flies (DF=1, 117; F=35.88; p<0.0001 for females; DF=1, 123; F=5.40; p=0.0218 for males) (D). Results consistent with these, but of weaker effect, were observed in flies derived from Mount Sinai (Figure S1). Error bars show 95% confidence intervals.

*InR^long^* allele was associated with greater lipid mass than *InR^short^* (Figures 5, S1); however, only alleles derived from Bowdoinham demonstrated a statistically significant difference when non-lipid mass was included as a covariate (p=0.0066 for females, p<0.0001 for males) (Table 1). Wings of *InR^long^* flies were also larger than those for *InR^short^*, at least in the Bowdoinham population. In that comparison, the average L2 wing vein length was 0.24 mm longer in *InR^long^* females (p<0.0001) and 0.12 mm longer in *InR^long^* males (p<0.0001) (Table 1, Figure 5). In flies derived from Mount Sinai, *InR^short^* females had the larger wings, but only by 0.02 mm (p=0.0029), and *InR^short^* and *InR^long^* males were not different from each other (p=0.5999) (Table 1, Figure S1).

An association between large size and *InR^long^*, the low-latitude allele, is predictable in terms of insulin signaling, but unpredictable given the observation that *Drosophila* are larger at higher latitude (Huey et al. 2000; De Jong & Bochdanovits 2003). Others have shown that proportionally larger wings relative to thorax size are associated with better flying in *Drosophila* (Hoffmann et al. 2007) and that the wing:thorax ratio increases with latitude (Karan et al. 1998), possibly because muscles in colder temperatures generate less power and proportionally larger wings can compensate for this (Gilchrist & Huey 2004). We measured thorax width to see if bigger wing:thorax ratios were associated with the high-latitude *InR^short^* allele, but they were not. *InR^short^* flies averaged smaller wing:thorax ratios relative to *InR^long^* flies (p<0.0001 for females; p=0.0218 for males) (Table 1, Figure 5). For all samples in which we measured the L2 wing vein, we also measured the L3 vein, and observed qualitatively identical results (data not shown).

***Stress tolerance.*** Reduction in IIS confers increased resistance to stress (Giannakou & Partridge 2007), and results for three of our four stress assays accord with these findings and our hypothesis that the high-latitude *InR^short^* allele mediates increased stress tolerance via reduced signaling (Figure 6). Females carrying the *InR^short^* allele recovered from chill coma on average 89 seconds faster than females carrying *InR^long^*; *InR^short^* males recovered 123 seconds faster (although this comparison was only statistically significant for males; p=0.1448 and p<0.0001, respectively) (Table 1, Figure 6). Likewise, *InR^short^* females and males exhibited 3.8 and 2.4 times the odds of surviving cold shock and 1.3 and 1.8 times the odds of surviving starvation, respectively, compared to *InR^long^* flies. However, the *InR^short^* allele did not correlate with higher tolerance of heat shock: instead, *InR^short^* flies showed 15.3 times the odds of dying in the assay compared to *InR^long^* flies (Table 2, Figure 6). All four of these outcomes are consistent with presumed selection pressures in natural populations, as flies in high latitude, temperate environments, where the *InR^short^* allele is common, likely face cold temperatures and scarce food availability during the winter, while flies in low latitude, semi-tropical environments, where *InR^long^* is common, should face thermal stress from higher temperatures. However, if levels of IIS mediate these stress phenotypes, the heat shock outcome provides a unique example of correlation reversal between signaling and stress. Nevertheless, the phenotypic correlations across these traits, and the correlations between the phenotypes and latitude, are patterns consistent with other examples of reciprocal tolerance for environmentally imposed thermal stresses (Hoffmann et al. 2001; Hoffmann et al. 2005).

**FIGURE 6.**
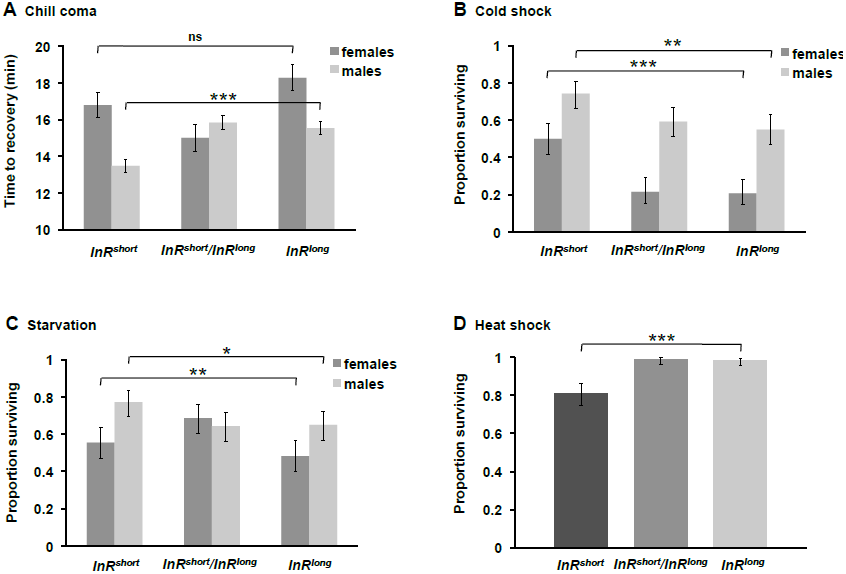
Average time to recovery from chill coma and proportion of flies surviving cold shock, starvation and heat shock. Although both *InR^short^* females and males recovered faster from chill coma than did *InR^long^* flies, the effect was only significant for males (DF=1, 187; F=2.14; p=0.1448 for females; DF=1, 207; F=16.31; p<0.0001) (A). A greater proportion of *InR^short^* flies survived cold shock (DF=2; χ^2^=35.70; p<0.0001 for females; DF=2; χ^2^=12.64; p=0.0018 for males) (B) and starvation (DF=2; χ^2^=12.36; p=0.0021 for females; DF=2; χ^2^=7.03; p=0.0298 for males) (C), but more *InR^long^* flies survived heat shock (DF=2; χ^2^=55.01; p<0.0001 for both sexes) (D). These flies were derived from the Mount Sinai population. Error bars show 95% confidence intervals.

**TABLE 2.** Statistical results for tests of temperature shock, starvation and lifespan for *InR* genotypes in a wild-type background.

| | Source | DF | $\chi^2$ | p |
| --- | --- | --- | --- | --- |
| Cold shock assay | genotype | 2 | 45.2572 | <0.0001 |
|  | sex | 1 | 93.799 | <0.0001 |
|  | genotype x sex | 2 | 3.0473 | 0.2179 |
| Starvation assay | genotype (f) | 2 | 12.3564 | 0.0021 |
|  | genotype (m) | 2 | 7.0283 | 0.0298 |
| Heat shock assay | genotype | 2 | 55.01 | <0.0001 |
|  | sex | 1 | 0.05 | 0.8294 |
|  | genotype x sex | 2 | 0.65 | 0.7228 |
| Lifespan | genotype (f) | 2 | 155.2047 | <0.0001 |
|  | genotype (m) | 2 | 165.8458 | <0.0001 |

***Lifespan.*** Insulin signaling has been established as a major regulator of longevity in many animals, including *Drosophila*: hypomorphic alleles of *InR* increase lifespan relative to wild-type controls (Tatar et al. 2001), as do other manipulations that reduce IIS (Clancy et al. 2001; Hwangbo et al. 2004; Grönke et al. 2010). Consequently, we hypothesized that *InR^short^* might increase lifespan relative to *InR^long^*. Flies carrying the *InR^short^* and *InR^long^* alleles showed little difference in rates of aging over the majority of lifespan, though *InR^short^* males exhibited reduced mortality late in life (Figure 7). The lifetime survivorship of *InR^short^* males was significantly longer than that of *InR^long^* males, with a mortality risk ratio of 1.3 for *InR^long^* over *InR^short^* (p=0.0095). The female mortality risk ratio for *InR^long^* over *InR^short^* was only 1.01, with no significant difference in longevity between these genotypes (p=0.8909). Interestingly, the heterozygote of both sexes lived significantly longer (p<0.0001) than either of the homozygote genotypes (Table 2). We hypothesize that the heterozygote longevity is a result of heterosis across the extended *InR*-embedded locus, a consequence of the fact that the homozygotes carried identical alleles with haplotypes extending beyond *InR*. Any general increase in fitness associated with greater heterozygosity was probably not mediated by IIS, as the heterozygotes were associated with intermediate levels of gene expression for the majority of IIS targets.

**FIGURE 7.**
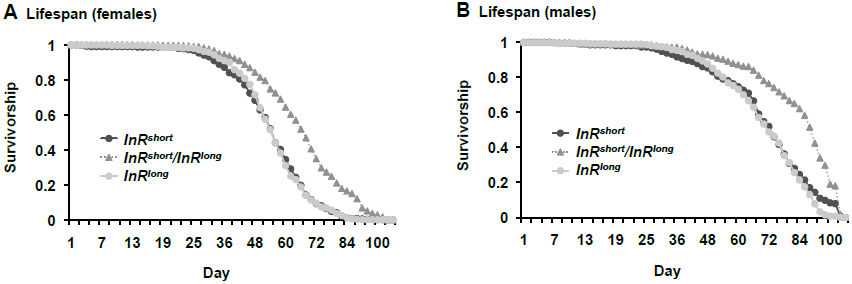
Survivorship curves, for females (A) and males (B), from flies derived from the Mount Sinai population. In both sexes the heterozygote lived significantly longer than either homozygote genotype (Table 2). In males, the *InR^short^* genotype exhibited a reduced rate of aging late in life to produce a significant lifespan extension relative to *InR^long^* (DF=1; χ^2^=6.73; p=0.0095), but there was no difference in longevity between *InR^short^* and *InR^long^* females (DF=1; χ^2^=0.02; p=0.8909).

***Allele effects in an* InR**-**background.** To examine how the *InR^short^* and *InR^long^* alleles performed in a genetic background with reduced InR activity, we crossed them into a stock with a hypomorphic allele (*InR^p5545^*) over a wild-type copy of *InR* on the *TM3* balancer chromosome (*InR^TM3^*), generating the four possible genotypic combinations. Previous work with this hypomorphic allele showed that a single copy of *InR^p5545^* reduces InR kinase activity by approximately half, and not as much as a double mutant (Tatar et al. 2001). The genotypic combinations allowed us to compare single copies of *InR^short^* to *InR^long^*, and wild-type flies to flies carrying a single copy of the hypomorphic *InR* mutant allele.

Consistent with the expectation that InR function increases fecundity, flies carrying the hypomorphic allele laid fewer eggs in the first 12 hours after mating than wild-type flies (p=0.0109). When paired with the hypomorphic allele, *InR^long^* females laid on average 38% more eggs than *InR^short^* females, which is consistent with the direct allele tests but was not statistically significant in this assay, and there was no difference in egg-laying between *InR^short^* and *InR^long^* in the wild-type background (Table S2, Figure 8). As in the direct allele tests on Bowdoinham flies, here patterns of development time were sex-specific. Females carrying the *InR* hypomorph developed 9.1 hours slower than *InR* wild-type flies (p<0.0001), but females carrying *InR^short^* and *InR^long^* were not different from each other. Males with *InR^short^* developed 1.3 hours slower than *InR^long^* males (p=0.0064), but males with or without the hypomorphic allele were not different from each other (Table S2, Figure 8). These patterns are consistent with the expected association of high InR activity and faster development, and the difference between the wild-type *InR^short^* and *InR^long^* alleles in males is much subtler than the difference between the wild-type and mutant alleles in females.

**FIGURE 8.**
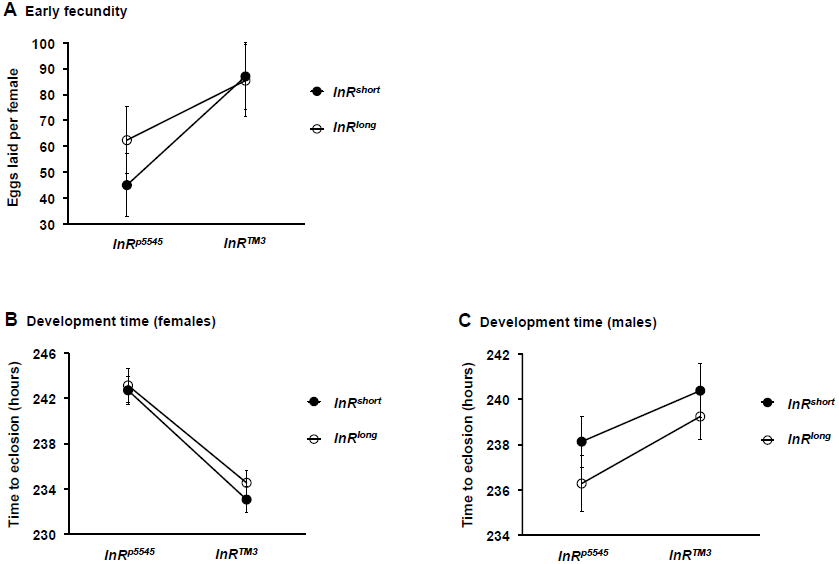
Average early fecundity (A) and development time (B, C) for single copies of *InR^short^* and *InR^long^*, paired with either an *InR* hypomorphic (*InR^p5545^*) or wild-type balancer (*InR^TM3^*) allele. Error bars show standard error of the mean.

Dry weight of wild-type flies was greater than those with the *InR* hypomorph (p=0.0061 for females; p=0.0763 for males), consistent with the prediction that increased *InR* activity results in larger mass, but there were no differences between *InR^short^* and *InR^long^* (Table S2, Figure 9). However, single copies of *InR^short^* and *InR^long^* were weakly associated with lesser and greater lipid levels, as we saw in the direct allele tests (p=0.0729 for females; p=0.0624 for males), but there were no differences between *InR* hypomorphic and wild-type alleles (Table S2, Figure 9). The observations of *InR^short^* with lower lipid level suggest that minor reduction in InR activity can reduce lipids even though the more general relationship between IIS and lipids is a negative one. This finding is in fact consistent with earlier characterization of *InR* mutant alleles, in which single copies of hypomorphic alleles were associated with reduced lipids (though not in our data), and double mutants with increased lipid content, relative to the wild-type (Tatar et al. 2001). Female wing sizes were not different from each other across any of the genotypic comparisons, but wings of *InR^long^* males were bigger than wings of *InR^short^* males (p=0.0179), as we saw in the direct allele tests (Table S2, Figure 10). The *InR^short^* and *InR^long^* alleles had no effect on the wing:thorax ratio, but both sexes had bigger ratios in the mutant background relative to the wild-type background (p<0.0001 for females; p=0.0001 for males) (Table S2, Figure 10). The correlation between the hypomorph and larger ratio suggests that decreased InR activity may increase the wing:thorax size relationship, which is what we observed in the direct allele tests as well.

**FIGURE 9.**
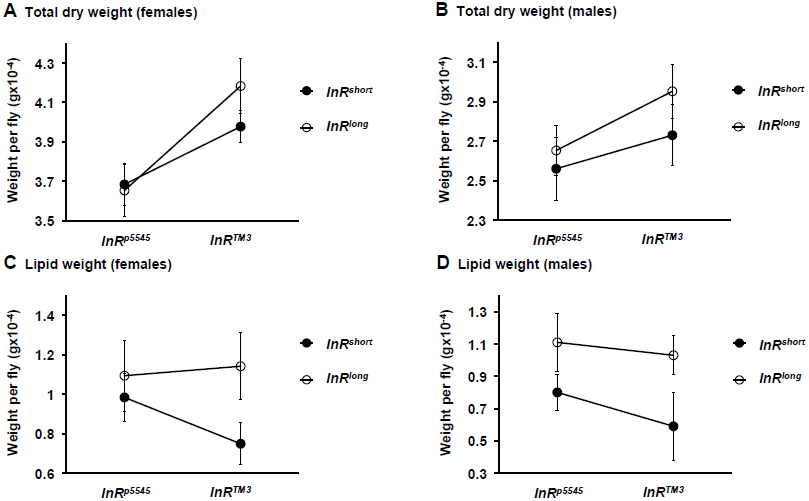
Average dry body weight (A, B) and average lipid weight (C, D) for single copies of *InR^short^* and *InR^long^*, paired with either an *InR* hypomorphic (*InR^p5545^*) or wild-type balancer (*InR^TM3^*) allele. Error bars show standard error of the mean.

**FIGURE 10.**
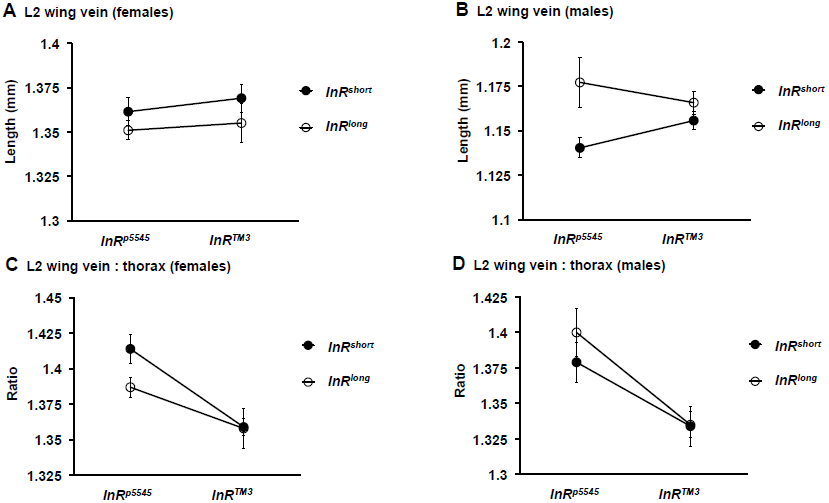
Average length of the L2 wing vein (A, B) and wing:thorax ratio (C, D) for single copies of *InR^short^* and *InR^long^*, paired with either an *InR* hypomorphic (*InR^p5545^*) or wild-type balancer (*InR*TM3**) allele. Error bars show standard error of the mean.

## DISCUSSION

Our results indicate that the complex indel polymorphism in the first exon of *InR* is likely a direct target of natural selection in wild populations of *D. melanogaster*. Allele frequencies exhibit persistent latitudinal clines, change rapidly and cyclically over seasonal timescales, and the two common alleles are pleiotropically associated with multiple traits that correspond to expected pressures in distinct climatic environments. The alleles are also associated with predicted up- and down-regulation of IIS, strongly suggesting that they mediate signaling behavior to effect differential trait expression. The fact that the alleles encode different amino acid sequences suggests that this occurs through changes in protein function. Overall our findings provide an example of life history tradeoffs influenced by a complex, strongly pleiotropic polymorphism.

While the phenotypes associated with *InR^short^* and *InR^long^* were specific and predictive, our assays did not precisely test the effects of the glutamine and histidine changes that account for the length differences in the complex indel polymorphism. This is for two reasons. First, the *InR^short^* and *InR^long^* alleles themselves contain several other SNPs, each of which exhibits nonrandom patterns of allele frequency across latitude and seasonal time (Figure 1). Only one of these (a non-synonymous SNP at position 17405614) has the potential to confound our functional results, however, as the other SNPs did not vary between our *InR^short^* and *InR^long^* lines. Allele frequencies at 17405614 exhibited substantial seasonal and clinal patterns, though not as strongly as several of the discrete indel variants that comprise the complex indel polymorphism (Figure 1). Second, genome regions flanking the complex indel polymorphsim were not fully randomized due to the limits of experimentally imposed recombination. Consequently, in our test lines, the *InR^short^* and *InR^long^* alleles were embedded in haplotypes that extended beyond the *InR* locus. Based on prior estimates of linkage disequilibrium, however, flanking variants should be unlinked to the focal indel polymorphism (Paaby et al. 2010). Although these other variants may be targets of selection—a conclusion supported by our observation that the entire *InR* locus exhibits elevated clinality and seasonality relative to the rest of the genome—the fact that the indel polymorphism exhibits such strong changes in allele frequency across climatic environments provides complementary evidence for allele functionality. This accords with other reports of high recombination and rapid decay of linkage disequilibrium (Mackay et al. 2012), and evidence that selection precisely targets functional loci, including *InR* (Fabian et al. 2012), in natural populations of *D. melanogaster*.

Two of the discrete polymorphisms associated with the complex indel polymorphism, 17405631 and 17405634-7, do not discriminate between *InR^short^* and *InR^long^* but rather discriminate between the published reference genome genotype (which is of intermediate amino acid length) and *InR^short^* or *InR^long^*. Both of these discrete polymorphisms show strong seasonal patterns of allele frequency, which suggests that the intermediate allele may be a target of selection as well. The discrete 17405631 and 17405634-7 alleles segregate at substantial frequencies in all populations, and in both North America and Australia the intermediate-length haplotype is the third most common allele class (Paaby et al. 2010). In our previous survey, the intermediate allele showed no cline in North America and a weak cline in Australia. Here, the discrete 17405631 and 17405634-7 alleles again show little evidence of clinality in North America (Figure 1D, Table S1). These observations support the conclusion that seasonal and geographical environments may impose different evolutionary forces, including aspects of demography and selection, even in the face of generally concordant responses to shared climatic selection pressure (Bergland et al. In Review).

Since flies derived from high latitudes live longer than those from low latitudes (Schmidt et al. 2005a), and reduction in IIS extends lifespan (Clancy et al. 2001; Tatar et al. 2001; Hwangbo et al. 2004; Grönke et al. 2010), we predicted that the *InR^short^* allele might be associated with longer lifespan than the *InR^long^* allele. *InR^short^* males did live significantly longer than *InR^long^* males, but the magnitude of difference was small compared to other traits and there were no differences between the alleles in females. The comparatively weak longevity effect may reflect decoupling between lifespan and reproduction that has been observed elsewhere (Khazaeli & Curtsinger 2010)—yet it may also represent true functional variation. Laboratory-derived *InR* mutations have shown a dramatic effect on lifespan, but only the heterozygous combination of two hypomorphic alleles, of all genotypes tested, significantly extended lifespan (Tatar et al. 2001). This suggests that IIS-mediated traits may be determined by precise titration of IIS levels.

In addition to longer male lifespan, the *InR^short^* allele was also associated with better cold and starvation tolerance and lower fecundity in females, a classic example of antagonistic pleiotropy (Williams 1957). The differences in fecundity were limited to early life, and represent variation in egg-laying rate, because total lifetime fecundity was not significantly different between the alleles. Modest differences in egg-laying in early life could have massive fitness consequences in the wild, particularly on patchy substrates with larval competition, yet the competitive advantage may be undetectable in the lab if fecundity is measured over more than one day. Consequently, we emphasize that estimates of reproduction and other quantitative fitness traits should be examined carefully, particularly in the context of decoupling correlated traits. We also note that although the high-latitude *InR^short^* allele is associated with smaller body size, in general high-latitude flies are bigger (Huey et al. 2000; De Jong & Bochdanovits 2003). If *InR* contributes to selection-mediated differences in body size, it either acts epistatically with other body-size loci or suffers antagonistic selection pressures across multiple fitness axes. This finding, like observations of the pleiotropic polymorphism at *neurofibromin 1* in which the high-latitude haplotype is negatively associated with wing size (Lee et al. 2013), demonstrates the complexity of selection dynamics in natural populations.

One hypothesis for strong genetic correlations among life history traits, including the tradeoff between lifespan and reproduction, is the presence of pleiotropic alleles. (Williams 1957; Flatt & Heyland 2011). If many genetic variants for life history are pleiotropic, then correlations among traits can never be completely dissolved. Our results implicate extensive pleiotropy, but the complexity of the *InR* indel polymorphism limits our ability to map phenotype to specific nucleotides. The two alleles we tested, *InR^short^* and *InR^long^*, differ at four amino acids across a span of 16 residues (Paaby et al. 2010). Even if in isolation these sites could break pleiotropy by functioning independently, over short timescales the distinction between pleiotropy and close linkage may be inconsequential (Paaby & Rockman 2013). However, this region at *InR* clearly experiences sufficient recombination (or mutation) over longer timescales to generate a hyper-variable set of alleles on which selection may act. Tests on additional *InR* alleles would help to resolve the functional roles of individual nucleotides. Moreover, our findings in support of pleiotropy may not represent the majority of life history alleles segregating in natural populations. Other work has shown that recombination can generate genotypes with positively correlated effects for both lifespan and reproduction, which suggests that life history variation in nature is likely determined by both pleiotropic and recombining non-pleiotropic loci (Khazaeli & Curtsinger 2013). Sampling genotypes from nature may represent blocks of alleles that produce the commonly observed correlations, but which can be broken under the right circumstances. Finally, although the two major alleles of the *InR* indel polymorphism showed canonical tradeoffs in the lab, the role of these alleles in the natural environment is unknown. Whether and how they effect trait expression might vary across different environments (Fournier-Level et al. 2013), with the possibility that under some conditions, pleiotropy may present as conditional neutrality (Anderson et al. 2011). Thus, true understanding of the role of this polymorphism in adaptive response, including fitness consequences and degree of pleiotropy, requires investigation rooted in the natural environment.

## ACKNOWLEDGEMENTS

We thank Katherine O'Brien for assistance in measuring the fly wings and Luke Noble for lending expertise on the pooled sample analyses. We also thank Li Yang and Nancy Bonini for sharing expertise and resources for qPCR. This work was supported by NSF DEB 0921307, NIH F32 GM097837 and the Charles H. Revson Foundation.

---

**FIGURE S1.**
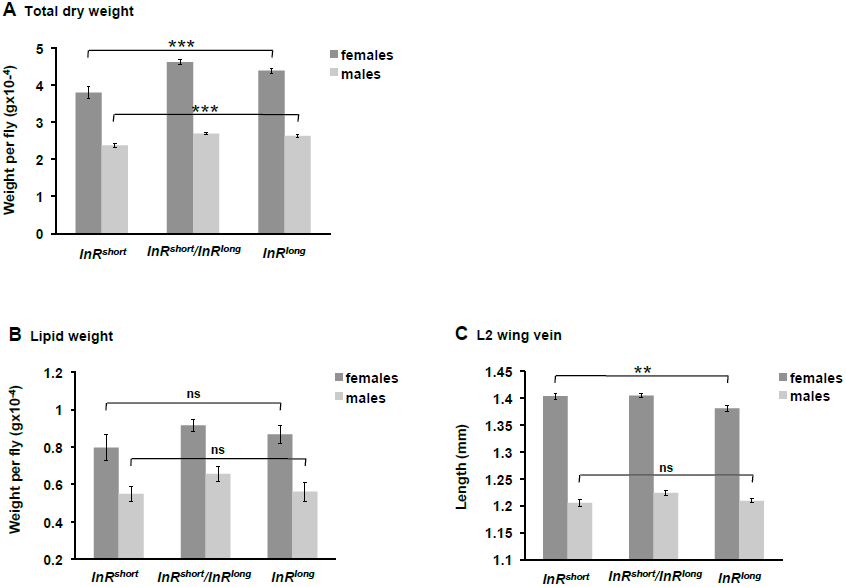
Average dry body weight, lipid weight, and length of the L2 wing vein for flies derived from the Mount Sinai population. *InRshort* flies had lower dry weight than *InRlong* flies (DF=1, 45; F=16.73; p=0.0002 for females; DF=1, 44; F=14.99; p=0.0004 for males) (A), but were not different in lipid weight (DF=1, 44; F=0.14; p=0.7140 for females; DF=1, 43; F=1.41; p=0.2414 for males) (B). *InRlong* females, but not males, had significantly larger wings than *InRshort* (DF= 1, 230; F=9.07; p=0.0029 for females; DF=1, 212; F=0.28; p=0.5999 for males) (C). Error bars show 95% confidence intervals.

**TABLE S1.** Characterization of the 8 discrete polymorphisms that are associated with, or are nearby, the complex insertion-deletion polymorphism and that are segregating at appreciable frequencies in the seasonal and clinal populations.

| Position on 3R | Discriminates between *InR<sup>short</sup>* and *InR<sup>long</sup>*? | Shows seasonality? | Shows clinality? |
| --- | --- | --- | --- |
| 17405614 | no | suggestive | yes |
| 17405619 | no | yes (low amplitude) | yes (low magnitude) |
| 17405631 | no | yes | no |
| 17405631-7 | yes | yes | weak |
| 17405634-7 | no | yes | weak |
| 17405637 | no | yes | weak |
| 17405651-4 | yes | suggestive | yes |
| 17405784 | yes | suggestive | yes |

**TABLE S2.** Statistical results for tests of *InR^short^* and *InR^long^* in an *InR*-background. Allele refers to *InR^short^* versus *InR^long^*; genotype refers to *InR^p5545^* versus *InR^TM3^*.

|  | Source | DF | SS | F | p |
| --- | --- | --- | --- | --- | --- |
| Early fecundity | allele | 1 | 1419.22 | 1.0665 | 0.3135 |
|  | genotype | 1 | 10378.41 | 7.7993 | 0.0109 |
|  | allele x genotype | 1 | 602.41 | 0.4527 | 0.5054 |
|  | line(allele) | 24 | 65966.5 | 2.0656 | 0.0487 |
| Development time | allele (f) | 1 | 124.0935 | 0.9696 | 0.3255 |
|  | genotype (f) | 1 | 8017.6243 | 62.6441 | <0.0001 |
|  | allele x genotype (f) | 1 | 11.0702 | 0.0865 | 0.7689 |
|  | line(allele) (f) | 19 | 5977.4714 | 2.4581 | 0.0008 |
|  | allele (m) | 1 | 658.434 | 7.5332 | 0.0064 |
|  | genotype (m) | 1 | 163.62 | 1.872 | 0.1722 |
|  | allele x genotype (m) | 1 | 150.351 | 1.7202 | 0.1906 |
|  | line(allele) (m) | 19 | 13405.72 | 8.0724 | <0.0001 |
| Dry weight | allele (f) | 1 | 0.00023442 | 0.3068 | 0.5891 |
|  | genotype (f) | 1 | 0.00816468 | 10.6846 | 0.0061 |
|  | allele x genotype (f) | 1 | 0.0000038 | 0.005 | 0.9448 |
|  | line(allele) (f) | 14 | 0.06566198 | 6.1377 | 0.0012 |
|  | allele (m) | 1 | 0.00001292 | 0.0036 | 0.9532 |
|  | genotype (m) | 1 | 0.01338558 | 3.7091 | 0.0763 |
|  | allele x genotype (m) | 1 | 0.00099214 | 0.2749 | 0.6089 |
|  | line(allele) (m) | 15 | 0.38324176 | 7.0798 | 0.0005 |
| Lipid weight | allele (f) | 1 | 0.0915883 | 3.8632 | 0.0729 |
|  | genotype (f) | 1 | 0.0142121 | 0.5595 | 0.4538 |
|  | allele x genotype (f) | 1 | 0.0709556 | 2.9929 | 0.1092 |
|  | line(allele) (f) | 14 | 1.1486353 | 3.4607 | 0.0187 |
|  | mass (f) | 1 | 0.0049022 | 0.2068 | 0.6574 |
|  | allele (m) | 1 | 0.096919 | 4.2567 | 0.0614 |
|  | genotype (m) | 1 | 0.0000176 | 0.0008 | 0.9783 |
|  | allele x genotype (m) | 1 | 0.0019569 | 0.0859 | 0.7744 |
|  | line(allele) (m) | 15 | 1.1553349 | 3.3828 | 0.0197 |
|  | mass (m) | 1 | 0.0187888 | 0.8252 | 0.3815 |
| L2 wing vein | allele (f) | 1 | 0.00000356 | 0.0022 | 0.963 |
|  | genotype (f) | 1 | 0.00156266 | 0.9506 | 0.3323 |
|  | allele x genotype (f) | 1 | 0.00003075 | 0.0187 | 0.8915 |
|  | line(allele) (f) | 11 | 0.03199929 | 1.7697 | 0.0717 |
|  | allele (m) | 1 | 0.00769703 | 5.8349 | 0.0179 |
|  | genotype (m) | 1 | 0.00006985 | 0.053 | 0.8186 |
|  | allele x genotype (m) | 1 | 0.00518159 | 3.928 | 0.0508 |
|  | line(allele) (m) | 10 | 0.0761658 | 5.7739 | <0.0001 |
| Wing:thorax ratio | allele (f) | 1 | 0.00024221 | 0.0896 | 0.7655 |
|  | genotype (f) | 1 | 0.04640658 | 17.1602 | <0.0001 |
|  | allele x genotype (f) | 1 | 0.00459448 | 1.6989 | 0.1961 |
|  | line(allele) (f) | 11 | 0.02335652 | 0.7852 | 0.6539 |
|  | allele (m) | 1 | 0.00398441 | 0.9221 | 0.3397 |
|  | genotype (m) | 1 | 0.069767 | 16.1468 | 0.0001 |
|  | allele x genotype (m) | 1 | 0.00321297 | 0.7436 | 0.391 |
|  | line(allele) (m) | 10 | 0.09228111 | 2.1357 | 0.0303 |

## LITERATURE CITED

Anderson, A. R., J. E. Collinge, A. A. Hoffmann, M. Kellett, and S. W. McKechnie. 2003. Thermal tolerance trade-offs associated with the right arm of chromosome 3 and marked by the *hsr-omega* gene in *Drosophila melanogaster*. Heredity 90:195–202.

Anderson, J. L., R. M. Reynolds, L. T. Morran, J. Tolman-Thompson, and P. C. Phillips. 2011. Experimental evolution reveals antagonistic pleiotropy in reproductive timing but not life span in *Caenorhabditis elegans*. J Gerontol A Biol Sci Med Sci 66:1300–1308.

Anderson, J. T., M. R. Wagner, C. A. Rushworth, K. V. Prasad, and T. Mitchell-Olds. 2013. The evolution of quantitative traits in complex environments. Heredity 112:4–12.

Anderson, J. T., J. H. Willis, and T. Mitchell-Olds. 2011. Evolutionary genetics of plant adaptation. Trends Genet 27:258–266.

Bansal, V. 2010. A statistical method for the detection of variants from next-generation resequencing of DNA pools. Bioinformatics 26:i318–324.

Bergland, A. O., H. S. Chae, Y. J. Kim, and M. Tatar. 2012. Fine-scale mapping of natural variation in fly fecundity identifies neuronal domain of expression and function of an aquaporin. PLoS Genet 8:e1002631.

Bergland, A. O., E. L. Behrman, K. R. O'Brien, P. S. Schmidt, and D. A. Petrov. In Review. Genomic evidence of rapid and stable adaptive oscillations over seasonal time scales in *Drosophila*.

Bock, I. R. and P. A. Parsons. 1981. Species of Australia and New Zealand. Pp. 291–308 *in* M. Ashburner, H. L. Carson, J. N. Thompson, ed. Genetics and Biology of Drosophila. Academic Press, London.

Capy, P., E. Pla, and J. R. David. 1993. Phenotypic and genetic variability of morphometrical traits in natural populations of *Drosophila melanogaster* and *Drosophila simulans*. Evolution 25:517–536.

Carbone, M. A., K. W. Jordan, R. F. Lyman, S. T. Harbison, J. Leips, T. J. Morgan, M. DeLuca, P. Awadalla, and T. F. Mackay. 2006. Phenotypic variation and natural selection at *catsup*, a pleiotropic quantitative trait gene in *Drosophila*. Curr Biol 16:912–919.

Clancy, D. J., D. Gems, L. G. Harshman, S. Oldham, H. Stocker, E. Hafen, S. J. Leevers, and L. Partridge. 2001. Extension of life-span by loss of CHICO, a *Drosophila* insulin receptor substrate protein. Science 292:104–106.

David, J. R. and C. Bocquet. 1975. Evolution in a cosmopolitan species: genetic latitudinal clines in *Drosophila melanogaster* wild populations. Experientia 31:164–166.

David, J. R. and P. Capy. 1988. Genetic variation of *Drosophila melanogaster* natural populations. Trends Genet 4:106–111.

De Jong, G. and Z. Bochdanovits. 2003. Latitudinal clines in *Drosophila melanogaster*: body size, allozyme frequencies, inversion frequencies, and the insulin-signalling pathway. J Genet 82:207–223.

De Luca, M., N. V. Roshina, G. L. Geiger-Thornsberry, R. F. Lyman, E. G. Pasyukova, and T. F. Mackay. 2003. *Dopa decarboxylase* (*Ddc*) affects variation in *Drosophila* longevity. Nat Genet 34:429–433.

Fabian, D. K., M. Kapun, V. Nolte, R. Kofler, P. S. Schmidt, C. Schlotterer, and T. Flatt. 2012. Genome-wide patterns of latitudinal differentiation among populations of *Drosophila melanogaster* from North America. Mol Ecol 21:4748–4769.

Flatt, T., K. J. Min, C. D'Alterio, E. Villa-Cuesta, J. Cumbers, R. Lehmann, D. L. Jones, and M. Tatar. 2008. *Drosophila* germ-line modulation of insulin signaling and lifespan. Proc Natl Acad Sci USA 105:6368–6373.

Flatt, T. and A. Heyland. 2011. Mechanisms of Life History Evolution: The Genetics and Physiology of Life History Traits and Trade-Offs. Oxford University Press, New York.

Fournier-Level, A., A. M. Wilczek, M. D. Cooper, J. L. Roe, J. Anderson, D. Eaton, B. T. Moyers, R. H. Petipas, R. N. Schaeffer, B. Pieper, M. Reymond, M. Koornneef, S. M. Welch, D. L. Remington, and J. Schmitt. 2013. Paths to selection on life history loci in different natural environments across the native range of *Arabidopsis thaliana*. Mol Ecol 22:3552–3566.

Giannakou, M. E. and L. Partridge. 2007. Role of insulin-like signalling in *Drosophila* lifespan. Trends Biochem Sci 32:180–188.

Gidaszewski, N. A., M. Baylac, and C. P. Klingenberg. 2009. Evolution of sexual dimorphism of wing shape in the *Drosophila melanogaster* subgroup. BMC Evol Biol 9:110.

Gilchrist, G. W. and R. B. Huey. 2004. Plastic and genetic variation in wing loading as a function of temperature within and among parallel clines in *Drosophila subobscura*. Integr Comp Biol 44:461–470.

Grönke, S., D. F. Clarke, S. Broughton, T. D. Andrews, and L. Partridge. 2010. Molecular evolution and functional characterization of *Drosophila* insulin-like peptides. PLoS Genet 6:e1000857.

Guirao-Rico, S. and M. Aguadé. 2009. Positive selection has driven the evolution of the *Drosophila Insulin-like receptor* (*InR*) at different timescales. Mol Biol Evol 26:1723–1732.

Harshman, L. G. and A. J. Zera. 2007. The cost of reproduction: the devil in the details. Trends Ecol Evol 22:80–86.

Hoffmann, A. A., R. Hallas, C. Sinclair, and P. Mitrovski. 2001. Levels of variation in stress resistance in *Drosophila* among strains, local populations, and geographic regions: patterns for desiccation, starvation, cold resistance, and associated traits. Evolution 55:1621–1630.

Hoffmann, A. A., E. Ratna, C. M. Sgrò, M. Barton, M. Blacket, R. Hallas, S. De Garis, and A. R. Weeks. 2007. Antagonistic selection between adult thorax and wing size in field released *Drosophila melanogaster* independent of thermal conditions. J Evol Biol 20:2219–2227.

Hoffmann, A. A. and A. R. Weeks. 2007. Climatic selection on genes and traits after a 100 year-old invasion: a critical look at the temperate-tropical clines in *Drosophila melanogaster* from eastern Australia. Genetica 129:133–147.

Hoffmann, A. A. and L. G. Harshman. 1999. Desiccation and starvation resistance in *Drosophila*: patterns of variation at the species, population and intrapopulation levels. Heredity 83:637–643.

Hoffmann, A. A., J. Shirriffs, and M. Scott. 2005. Relative importance of plastic vs genetic factors in adaptive differentiation: geographical variation for stress resistance in *Drosophila melanogaster* from eastern Australia. Functional Ecology 19:222–227.

Huey, R. B., G. W. Gilchrist, M. L. Carlson, D. Berrigan, and L. Serra. 2000. Rapid evolution of a geographic cline in size in an introduced fly. Science 287:308–309.

Hwangbo, D. S., B. Gershman, M. P. Tu, M. Palmer, and M. Tatar. 2004. Drosophila dFOXO controls lifespan and regulates insulin signalling in brain and fat body. Nature 429:562–566.

Jones, F. C., M. G. Grabherr, Y. F. Chan, P. Russell, E. Mauceli, J. Johnson, R. Swofford, M. Pirun, M. C. Zody, S. White, E. Birney, S. Searle, J. Schmutz, J. Grimwood, M. C. Dickson, R. M. Myers, C. T. Miller, B. R. Summers, A. K. Knecht, S. D. Brady, H. Zhang, A. A. Pollen, T. Howes, C. Amemiya, J. Baldwin, T. Bloom, D. B. Jaffe, R. Nicol, J. Wilkinson, E. S. Lander, F. Di Palma, K. Lindblad-Toh, and D. M. Kingsley. 2012. The genomic basis of adaptive evolution in threespine sticklebacks. Nature 484:55–61.

Karan, D., A. K. Munjal, P. Gibert, B. Moreteau, R. Parkash, and J. R. David. 1998. Latitudinal clines for morphometrical traits in *Drosophila kikkawai*: a study of natural populations from the Indian subcontinent. Genet Res 71:31–38.

Khazaeli, A. A. and J. W. Curtsinger. 2010. Life history variation in an artificially selected population of *Drosophila melanogaster*: pleiotropy, superflies, and age-specific adaptation. Evolution 64:3409–3416.

Khazaeli, A. A. and J. W. Curtsinger. 2013. Pleiotropy and life history evolution in *Drosophila melanogaster*: uncoupling life span and early fecundity. J Gerontol A Biol Sci Med Sci 68:546–553.

Lee, S. F., Y. C. Eyre-Walker, R. V. Rane, C. Reuter, G. Vinti, L. Rako, L. Partridge, and A. A. Hoffmann. 2013. Polymorphism in the neurofibromin gene, *Nf1*, is associated with antagonistic selection on wing size and development time in *Drosophila melanogaster*. Mol Ecol 22:2716–2725.

Leroi, A. M., A. Bartke, G. De Benedictis, C. Franceschi, A. Gartner, E. S. Gonos, M. E. Fedei, T. Kivisild, S. Lee, N. Kartaf-Ozer, M. Schumacher, E. Sikora, E. Slagboom, M. Tatar, A. I. Yashin, J. Vijg, and B. Zwaan. 2005. What evidence is there for the existence of individual genes with antagonistic pleiotropic effects? Mech Ageing Dev 126:421–429.

Li, H. and R. Durbin. 2010. Fast and accurate long-read alignment with Burrows-Wheeler transform. Bioinformatics 26:589–595.

Linnen, C. R., Y. P. Poh, B. K. Peterson, R. D. Barrett, J. G. Larson, J. D. Jensen, and H. E. Hoekstra. 2013. Adaptive evolution of multiple traits through multiple mutations at a single gene. Science 339:1312–1316.

Mackay, T. F., S. Richards, E. A. Stone, A. Barbadilla, J. F. Ayroles, D. Zhu, S. Casillas, Y. Han, M. M. Magwire, J. M. Cridland, M. F. Richardson, R. R. Anholt, M. Barron, C. Bess, K. P. Blankenburg, M. A. Carbone, D. Castellano, L. Chaboub, L. Duncan, Z. Harris, M. Javaid, J. C. Jayaseelan, S. N. Jhangiani, K. W. Jordan, F. Lara, F. Lawrence, S. L. Lee, P. Librado, R. S. Linheiro, R. F. Lyman, A. J. Mackey, M. Munidasa, D. M. Muzny, L. Nazareth, I. Newsham, L. Perales, L. L. Pu, C. Qu, M. Ramia, J. G. Reid, S. M. Rollmann, J. Rozas, N. Saada, L. Turlapati, K. C. Worley, Y. Q. Wu, A. Yamamoto, Y. Zhu, C. M. Bergman, K. R. Thornton, D. Mittelman, and R. A. Gibbs. 2012. The *Drosophila melanogaster* Genetic Reference Panel. Nature 482:173–178.

McKenna, A., M. Hanna, E. Banks, A. Sivachenko, K. Cibulskis, A. Kernytsky, K. Garimella, D. Altshuler, S. Gabriel, M. Daly, and M. A. DePristo. 2010. The Genome Analysis Toolkit: a MapReduce framework for analyzing next-generation DNA sequencing data. Genome Res 20:1297–1303.

Méndez-Vigo, B., J. M. Martínez-Zapater, and C. Alonso-Blanco. 2013. The flowering repressor *SVP* underlies a novel *Arabidopsis thaliana* QTL interacting with the genetic background. PLoS Genet 9:e1003289.

Mitrovski, P. and A. A. Hoffmann. 2001. Postponed reproduction as an adaptation to winter conditions in *Drosophila melanogaster*: evidence for clinal variation under semi-natural conditions. Proc Biol Sci 268:2163–2168.

Paaby, A. B., M. J. Blacket, A. A. Hoffmann, and P. S. Schmidt. 2010. Identification of a candidate adaptive polymorphism for *Drosophila* life history by parallel independent clines on two continents. Mol Ecol 19:760–774.

Paaby, A. B. and M. V. Rockman. 2013. The many faces of pleiotropy. Trends Genet 29:66–73.

Paaby, A. B. and P. S. Schmidt. 2008. Functional significance of allelic variation at *methuselah*, an aging gene in *Drosophila*. PloS One 3:e1987.

Paaby, A. B. and P. S. Schmidt. 2009. Dissecting the genetics of longevity in *Drosophila melanogaster*. Fly 3:29–38.

Partridge, L. and D. Gems. 2002. The evolution of longevity. Curr Biol 12:R544–546.

Partridge, L., D. Gems, and D. J. Withers. 2005. Sex and death: what is the connection? Cell 120:461–472.

Partridge, L., N. Prowse, and P. Pignatelli. 1999. Another set of responses and correlated responses to selection on age at reproduction in *Drosophila melanogaster*. Proc Biol Sci 266:255–261.

Rako, L., M. J. Blacket, S. W. McKechnie, and A. A. Hoffmann. 2007. Candidate genes and thermal phenotypes: identifying ecologically important genetic variation for thermotolerance in the Australian *Drosophila melanogaster* cline. Mol Ecol 16:2948–2957.

Remolina, S. C., P. L. Chang, J. Leips, S. V. Nuzhdin, and K. A. Hughes. 2012. Genomic basis of aging and life-history evolution in *Drosophila melanogaster*. Evolution 66:3390–3403.

Reznick, D. 1985. Costs of reproduction: an evaluation of the empirical evidence. Oikos 44:257–267.

Rohlf, F. J. 2008. TpsDIG, State University of New York at Stony Brook.

Schmidt, P. S., L. Matzkin, M. Ippolito, and W. F. Eanes. 2005a. Geographic variation in diapause incidence, life-history traits, and climatic adaptation in *Drosophila melanogaster*. Evolution 59:1721–1732.

Schmidt, P. S., A. B. Paaby, and M. S. Heschel. 2005b. Genetic variance for diapause expression and associated life histories in *Drosophila melanogaster*. Evolution 59:2616–2625.

Schmidt, P. S. and A. B. Paaby. 2008. Reproductive diapause and life-history clines in North American populations of *Drosophila melanogaster*. Evolution 62:1204–1215.

Schmidt, P. S., C. T. Zhu, J. Das, M. Batavia, L. Yang, and W. F. Eanes. 2008. An amino acid polymorphism in the *couch potato* gene forms the basis for climatic adaptation in *Drosophila melanogaster*. Proc Natl Acad Sci USA 105:16207–16211.

Sgrò, C. M., B. van Heerwaarden, V. Kellermann, C. W. Wee, A. A. Hoffmann, and S. F. Lee. 2013. Complexity of the genetic basis of ageing in nature revealed by a clinal study of lifespan and *methuselah*, a gene for ageing, in *Drosophila* from eastern Australia. Mol Ecol 22:3539–3551.

Stearns, S. C. 1991. Trade-offs in life-history evolution. Functional Ecology 3:259–268.

Stearns, S. C. 1992. The Evolution of Life Histories. Oxford University Press, New York.

Tatar, M., A. Bartke, and A. Antebi. 2003. The endocrine regulation of aging by insulin-like signals. Science 299:1346–1351.

Tatar, M., A. Kopelman, D. Epstein, M. P. Tu, C. M. Yin, and R. S. Garofalo. 2001. A mutant *Drosophila* insulin receptor homolog that extends life-span and impairs neuroendocrine function. Science 292:107–110.

Toivonen, J. M. and L. Partridge. 2009. Endocrine regulation of aging and reproduction in *Drosophila*. Mol Cell Endocrinol 299:39–50.

Trotta, V., F. C. Calboli, M. Ziosi, D. Guerra, M. C. Pezzoli, J. R. David, and S. Cavicchi. 2006. Thermal plasticity in *Drosophila melanogaster*: a comparison of geographic populations. BMC Evol Biol 6:67.

Vermeulen, C. J. and V. Loeschcke. 2007. Longevity and the stress response in *Drosophila*. Exp Geront 42:153–159.

Williams, G. C. 1957. Pleiotropy, natural selection, and the evolution of senescence. Evolution 11:398–411.

